## Supplementary information for "Development of an improved blood-stage malaria vaccine targeting the essential RH5-CyRPA-RIPR invasion complex"

#### Supplementary Table and Figure legends

**Table S1 – Details of all monoclonal antibodies used in this study.**
NA = Not applicable. NT = Not tested. RH5 epitope bin refers to colour coding reported by Alanine *et al*[1].

**Table S2 – Summary statistics for rat immunisation study using full-length RCR antigens.**
Summary statistics for ELISA data shown in **Figure 4A-C**. Sections A-C of the table correspond to **Figure 4A-C**. Significance determined by one-way ANOVA with Dunnett's multiple comparisons test versus the single antigen only group as indicated, * $p$<0.05, *** $p$<0.0001, **** $p$<0.0001, ns = not significant.

**Table S3 – Summary statistics for rat immunisation study using R58C and R78C antigens.**
Summary statistics for ELISA data shown in **Figure 6A-C**. Sections A-C of the table correspond to **Figure 6A-C**. Significance determined by one-way ANOVA of data with Dunnett's multiple comparisons test versus the single antigen only group as indicated, * $p$<0.05, ** $p$<0.001, **** $p$<0.0001, ns = not significant.

**Table S4 – Details of primers used to generate RIPR truncations**

**Table S5 – Details of RIPR truncations expressed**

**Table S6 – Amino acid sequences of R78C and R58C**

**Figure S1 – GIA of anti-RH5, -CyRPA, and -RIPR monoclonal antibodies and representative data for mAb interactions with the RCR-complex.**
Single cycle GIA assays were performed using *P. falciparum* 3D7 clone parasites. Values reported are analysis at 0.5 mg/mL test mAb concentration unless otherwise stated. (**A**) Anti-RH5 mAbs, colour coded as per epitope bins reported previously[1] and as **Figure 2A**. 2AC7 and 9AD4 were tested at 0.2 mg/mL. (**B**) Anti-CyRPA mAbs colour coded as per epitope bins reported previously[2]. (**C**) Anti-RIPR mAbs colour coded as per **Figure 5A**. Each point is the mean of three technical repeats. Bars show mean values. GIA below the line at 20 % is regarded as negative. (**D**) Size exclusion chromatograms showing complex formation analysis between the RCR-complex and anti-RH5 mAb R5.011 [Type I, GIA-negative] and non-reducing SDS-PAGE gel of co-immunoprecipitation of the pre-formed recombinant RCR-complex without (-) or with (+) mAb R5.011 bound to protein G agarose beads. Other representative examples shown for (**E**) anti-CyRPA mAb Cy.009 [Type I; GIA-positive]; (**F**) anti-CyRPA mAb Cy.002 [Type II; GIA-negative]; (**G**) anti-RIPR mAb RP.012 [Type I; GIA-positive]; and (**H**) anti-RIPR mAb RP.017 [Type II; GIA-negative].

**Figure S2 – Analysis of inter-antigen synergistic GIA of monoclonal antibodies targeting the RCR-complex.**
Full GIA dilution curves of pairwise mAb combinations targeting the RCR-complex. Black: Titrated mAb alone. Red: predicted Bliss additivity GIA for combination of the two mAbs with one held constant at ~30 % GIA. Blue: measured GIA data of the combination of the titrated mAb plus held concentration of the other mAb. Individual points are the mean of a triplicate measurement with error bars indicating the standard deviation.

**Figure S3 – Rat immunisation study with full-length RCR-complex antigen combinations.**
Wistar rats were immunised intramuscularly with soluble protein vaccines formulated in Matrix-M™ adjuvant on days 0, 28 and 56. Tail bleeds were taken on day -2 (pre-immunisation) and day 42 (post-second dose), followed by terminal bleed on day 70 (post-third dose). (**A**) Serum IgG ELISA data shown for samples taken on days 42 and 70 against full-length RH5 (red), CyRPA (blue), and RIPR (green) reported in µg/mL. Individual and median group responses are shown. Dotted line corresponds to median antigen-specific IgG response from the relevant group of single antigen immunised animals. (**B**) Single-cycle GIA assay using *P. falciparum* clone 3D7. Total IgG was purified from day 70 serum samples and titrated in the GIA assay. Each line represents an individual animal, and each point represents the mean of three technical replicates. GIA at 1 mg/mL total purified IgG was interpolated for each animal. (**C**) Data from (B) replotted against total antigen-specific IgG concentration in µg/mL as measured by ELISA in each purified total IgG sample. Each dataset was fitted with a Richard's five-parameter dose-response curve with no constraints to ascertain the $EC_{50}$. Doses of each antigen in µg used for vaccination are specified in brackets in the title of each graph.

**Figure S4 – Calibration free concentration analysis (CFCA) for rat IgG and rat immunisation study with the single full-length RH5, CyRPA and RIPR antigens.**
(**A**) CFCA was performed using a Biacore X100 instrument to measure absolute (µg/mL) concentrations of antigen-specific antibody in purified IgG samples from vaccinated rats. Correlation of anti-RH5 IgG (shown in red) measured in arbitrary units (AU) by ELISA for each sample and the RH5-specific IgG concentration measured by CFCA. Linear regression $r^2$ value is shown (N=6); with the slope used to define the conversion factor between ELISA AU and antigen-specific IgG concentration in ng/mL. The same analysis was repeated for CyRPA (N=6; blue), and RIPR (N=10; green). (**B**) Correlation of total antigen-specific IgG titers from purified IgG measured using the R+C+R standardised ELISA in arbitrary units (AU) versus the summed total antigen-specific responses in µg/mL (derived from independent measurement of responses to the three individual antigens by standardised ELISAs converted to µg/mL by CFCA). Significance determined by Pearson's correlation coefficient (r). (**C**) Correlation of interpolated total antigen-specific IgG $EC_{50}$ values in the GIA assay derived using the R+C+R ELISA in AU or the summed µg/mL. Significance determined by Pearson's correlation coefficient (r). (**D-F**) Serum from Wistar rats immunised three times with full-length antigens formulated in Matrix-M™ adjuvant were tested by ELISA on days -2, 42 and 70 for anti-antigen total IgG responses (reported in µg/mL). (**D**) Immunisation with 0.2 µg (hollow circles) or 2 µg (filled circles) RH5. (**E**) Immunisation with 2 µg (hollow circles) or 20 µg (filled circles) CyRPA. (**F**) Immunisation with 2 µg (hollow circles) or 20 µg (filled circles) RIPR. (**G-J**) Single cycle GIA assay against *P. falciparum* 3D7 clone using total IgG purified from serum of (**G**) naïve and Matrix-M™ only immunised rats, (**H**) RH5, (**I**) CyRPA, and (**J**) RIPR immunised animals. (**K-M**) Functional quality analysis of GIA data from immunisations with full-length RH5, CyRPA or RIPR. Full GIA curves from

(**H-J**) re-plotted against the appropriate anti-antigen specific IgG content of each sample as measured by ELISA in µg/mL. N=12 animals per antigen, each dataset fitted with a Richard's five-parameter dose-response curve with no constraints. The dashed line shows 0 % GIA and dotted line shows 50 % GIA for comparison. (**N**) Summary table of interpolated GIA $EC_{50}$ results for each immunisation antigen.

**Figure S5 – Production of RIPR protein fragments, antigen-reversal GIA assays and anti-RIPR mAb epitope mapping.**
(**A**) Schematic of full-length RIPR protein (RIPR FL) with the design of each RIPR protein fragment labelled with first and last amino acid. EGF domains shown as blue circles. Chen3 and Chen1/2 were previously reported as "Ripr3" and "Ripr1/2" by Chen *et al.*[3] (**B**) Reducing SDS-PAGE of RIPR FL and RIPR fragments fused to the monoFc prior to cleavage by TEV protease. CTD = C-terminal domain. MonoFc-GFP fusion protein produced as a control. (**C**) Single cycle GIA assay using *P. falciparum* clone 3D7 of purified IgG from rabbits immunised with full-length RIPR (blue, N=6) or from a pre-immune serum negative control (grey, N=1). Each dataset fitted with a Richard's five-parameter dose-response curve with no constraints. Mean of triplicate wells and SD shown. (**D**) Antigen reversal single cycle GIA assay using *P. falciparum* clone 3D7 and pooled anti-RIPR IgG from (C). Total purified IgG was held at 3 mg/mL and the indicated RIPR proteins were included at 1 µM final concentration unless otherwise indicated. Blue bar = anti-RIPR IgG alone; white bars = anti-RIPR IgG + indicated protein; grey bars = indicated protein alone; green bars = anti-RH5 polyclonal IgG control (pAb) with (+) or without (-) addition of RIPR EGF(7-8) protein. Mean and SD of triplicate wells shown. Significance determined by Kruskal-Wallis test with Dunn's multiple comparison post-test versus anti-RIPR FL IgG alone; * $p<0.05$, ** $p<0.01$. (**E**) Antigen reversal single cycle GIA assay using *P. falciparum* clone 3D7 and pooled anti-RIPR FL IgG from (C). Total purified IgG was held at 3 mg/mL and the indicated RIPR EGF proteins were titrated from 20 µM to 0.31 µM final concentration. Coloured bars = anti-RIPR IgG + indicated protein; grey bars = indicated protein alone; blue bars = controls: Control 1 = 3 mg/mL anti-RIPR IgG alone. Control 2 = 20 µM RIPR EGF(5-6) or RIPR EGF(7-8) alone. Control 3 = anti-RH5 IgG at 3 mg/mL and Control 4 = 20 µM indicated RIPR EGF protein with anti-RH5 IgG at 3 mg/mL. Control 5 = RIPR EGF(5-6) and RIPR EGF(7-8) proteins mixed at 20 µM and tested alone. All bars are mean and error bars are SD, N=3 except for EGF(5-8) where N=6. (**F**) Identification of anti-RIPR mAb binding regions by dot blot using full-length RIPR and indicated RIPR protein fragments. A subset of mAbs were also tested against denatured full-length RIPR protein. (**G**) Anti-RIPR mAbs RP.016 and RP.017 bound denatured RIPR and Nhalf by dot blot and were then screened on an overlapping peptide ELISA covering amino acids 21-247 and 364-648; both mAbs bind to peptide 1 (IDLIEGIFYEKNEIDKLTFS). Representative graph of two repeats shown.

**Figure S6 – Investigation of RIPR interactions with SEMA7A and PTRAMP-CSS.**
(**A**) SPR sensorgrams show multi-cycle kinetics of CyRPA (left) or SEMA7A (right) binding to RIPR coated on a CM5 chip. Report points of the equilibrium binding levels of the 9-step 2-fold dilutions for CyRPA binding to RIPR are fitted to the equilibrium or steady-state binding model for $K_D$ determination (middle). No RIPR binding detected for SEMA7A. (**B**) SPR sensorgrams show multi-cycle kinetics of MTRAP (left) or RIPR (right) binding to SEMA7A coated on a CM5 chip. Report points of the equilibrium binding levels of the 9-step 2-fold dilutions for MTRAP binding to SEMA7A are fitted to the equilibrium or steady-state binding model for $K_D$ determination (middle). No SEMA7A binding detected for RIPR. (**C**) SDS-PAGE gel showing the recombinant PTRAMP-CSS heterodimer run

under non-reducing (N) and reducing (R) conditions. Arrows indicate PTRAMP-CSS heterodimer (expected mass 62 kDa, PTRAMP and CSS 31 kDa each), confirmed by mass spectrometry analysis (data not shown). Higher band is an unidentified contaminant from the baculovirus expression system. (**D**) Western blot of gel shown in (C) using streptavidin-AP to detect biotin added to the BAP tag by BirA. (**E**) SPR sensorgrams show multi-cycle kinetics of RIPR binding to PTRAMP-CSS heterodimer coated on a CM5 chip. Curves show a 5-step 2-fold dilution series for RIPR binding. The equilibrium or steady-state binding model was used for $K_D$ determination. (**F**) No binding of RIPR EGF(5-8) to the PTRAMP-CSS heterodimer could be detected by the same method as (E).

**Figure S7 – Development of vaccines containing RIPR EGF-like domains.**
(**A**) Wistar rats were immunised three times intramuscularly on days 0, 28 and 56 with full-length (FL) RIPR or the indicated EGF domains as soluble protein vaccines, or with HBsAg VLPs conjugated to the indicated EGF domains or HBsAg VLP control, all formulated in 25 µg Matrix-M™ adjuvant. All doses given as equimolar to 2 µg full-length RIPR. Tail bleeds were taken on day -2 (pre-immunisation), day 27 (post first dose), and a terminal bleed was taken on day 55 (post-second dose). Serum IgG ELISA data shown against full-length RIPR (green) reported in µg/mL. Individual and median group responses are shown. (**B**) Structural model of R78C. Blue: CyRPA (PDB: 5TIK[4]); orange: linker and SpyTag; green: RIPR EGF(7-8) (AlphaFold[5]). (**C**) Structural model of R58C. Blue: CyRPA (PDB: 5TIK[4]), orange: linker and SpyTag, green: RIPR EGF(5-8) (AlphaFold[5]). (**D**) Reducing SDS-PAGE gel of full-length antigens (predicted molecular weights: RH5: 65 kDa, CyRPA: 42 kDa, RIPR 120 kDa), R78C (54 kDa) and R58C (66 kDa). (**E**) ELISA of a selection of GIA-positive anti-CyRPA, -RIPR, and -RH5 mAbs against R78C and R58C proteins. Error bars are SD (3 technical replicates), dotted line is the negative cut off. (**F**) Antigen reversal GIA assay using R58C protein and pooled anti-RIPR full-length rabbit IgG from **Figure S5C**, and (**G**) pooled anti-CyRPA rat IgG from **Figure S4E**. Total purified IgG (polyclonal antibody, pAb) was held at the indicated mg/mL concentration and R58C protein was titrated from 20 µM to 0.31 µM final concentration. Coloured bars = anti-RIPR IgG + R58C protein (green) or anti-CyRPA IgG + R58C protein (blue). Grey bars show indicated protein tested alone, or pAb control, or anti-RH5 pAb IgG negative control with or without 20 µM R58C protein. (**H**) Size exclusion chromatograms demonstrating complex formation between RH5 and R78C. (**I**) Non-reducing SDS-PAGE gel assessing binary complex formation between RH5 and R78C. Asterisks on chromatograms indicate which gel lanes correspond to the peaks in (H).

**Figure S8 – Rat immunisation study with R78C and R58C vaccines.**
Wistar rats were immunised intramuscularly with soluble protein vaccines formulated in 25 µg Matrix-M™ adjuvant on days 0, 28 and 56. Tail bleeds were taken on day -2 (pre-immunisation), day 14 (post-first dose) and day 42 (post-second dose), followed by terminal bleed on day 70 (post-third dose). (**A**) Serum IgG ELISA data shown for samples taken on days 14, 42 and 70 against full-length RH5 (red), CyRPA (blue), and RIPR (green) reported in µg/mL. Individual and median group responses are shown. Dotted line corresponds to median antigen-specific IgG response from the relevant group of single antigen immunised animals where applicable. Data are spread over two graphs to show separate immunisation cohorts. (**B**) Single-cycle GIA assay using *P. falciparum* clone 3D7. Total IgG was purified from day 70 serum samples and titrated in the GIA assay. Each line represents an individual animal, and each point represents the mean of three technical replicates. GIA at 1 mg/mL total purified IgG was interpolated for each animal. (**C**) Data from (B) replotted against total antigen-specific IgG concentration in µg/mL as measured by ELISA in each purified total IgG sample. Each

dataset was fitted with a Richard's five-parameter dose-response curve with no constraints to ascertain the $EC_{50}$. Doses in µg of each antigen used for vaccination are specified in brackets in the title on each graph.

#### Supplementary information references

Figure S1

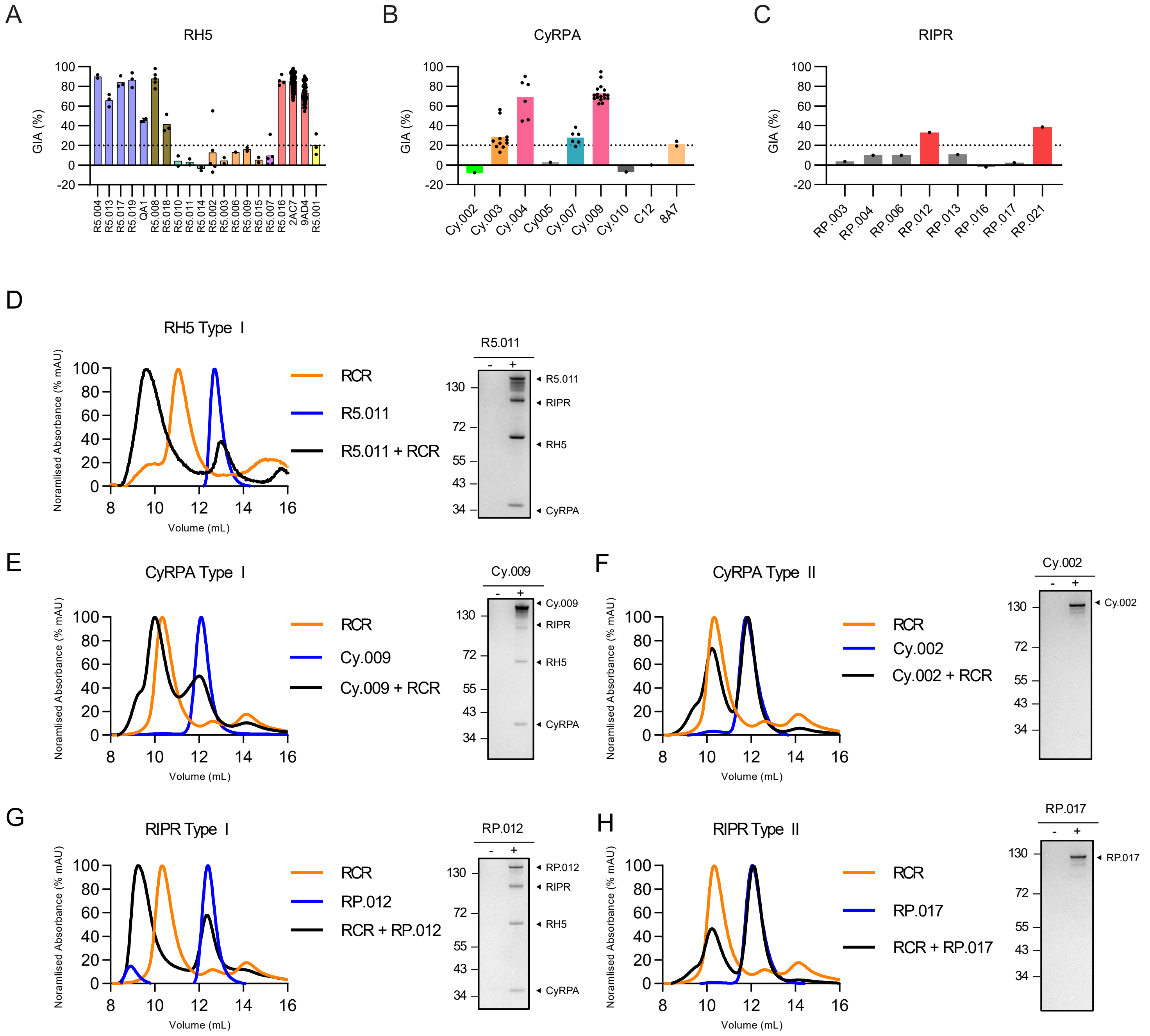

A
RH5
GIA (%)
R5.004
R5.013
R5.017
R5.019
QA1
R5.008
R5.018
R5.010
R5.011
R5.014
R5.002
R5.003
R5.006
R5.009
R5.015
R5.007
R5.016
2AC7
9AD4
R5.001
B
CyRPA
GIA (%)
Cy.002
Cy.003
Cy.004
Cy005
Cy.007
Cy.009
Cy.010
C12
8A7
C
RIPR
GIA (%)
RP.003
RP.004
RP.006
RP.012
RP.013
RP.016
RP.017
RP.021
D
RH5 Type I
Noramlised Absorbance (% mAU)
Volume (mL)
RCR
R5.011
R5.011 + RCR
R5.011
R5.011
RIPR
RH5
CyRPA
E
CyRPA Type I
Noramlised Absorbance (% mAU)
Volume (mL)
RCR
Cy.009
Cy.009 + RCR
Cy.009
Cy.009
RIPR
RH5
CyRPA
F
CyRPA Type II
Noramlised Absorbance (% mAU)
Volume (mL)
RCR
Cy.002
Cy.002 + RCR
Cy.002
Cy.002
G
RIPR Type I
Noramlised Absorbance (% mAU)
Volume (mL)
RCR
RP.012
RCR + RP.012
RP.012
RP.012
RIPR
RH5
CyRPA
H
RIPR Type II
Noramlised Absorbance (% mAU)
Volume (mL)
RCR
RP.017
RCR + RP.017
RP.017
RP.017

### Figure S2

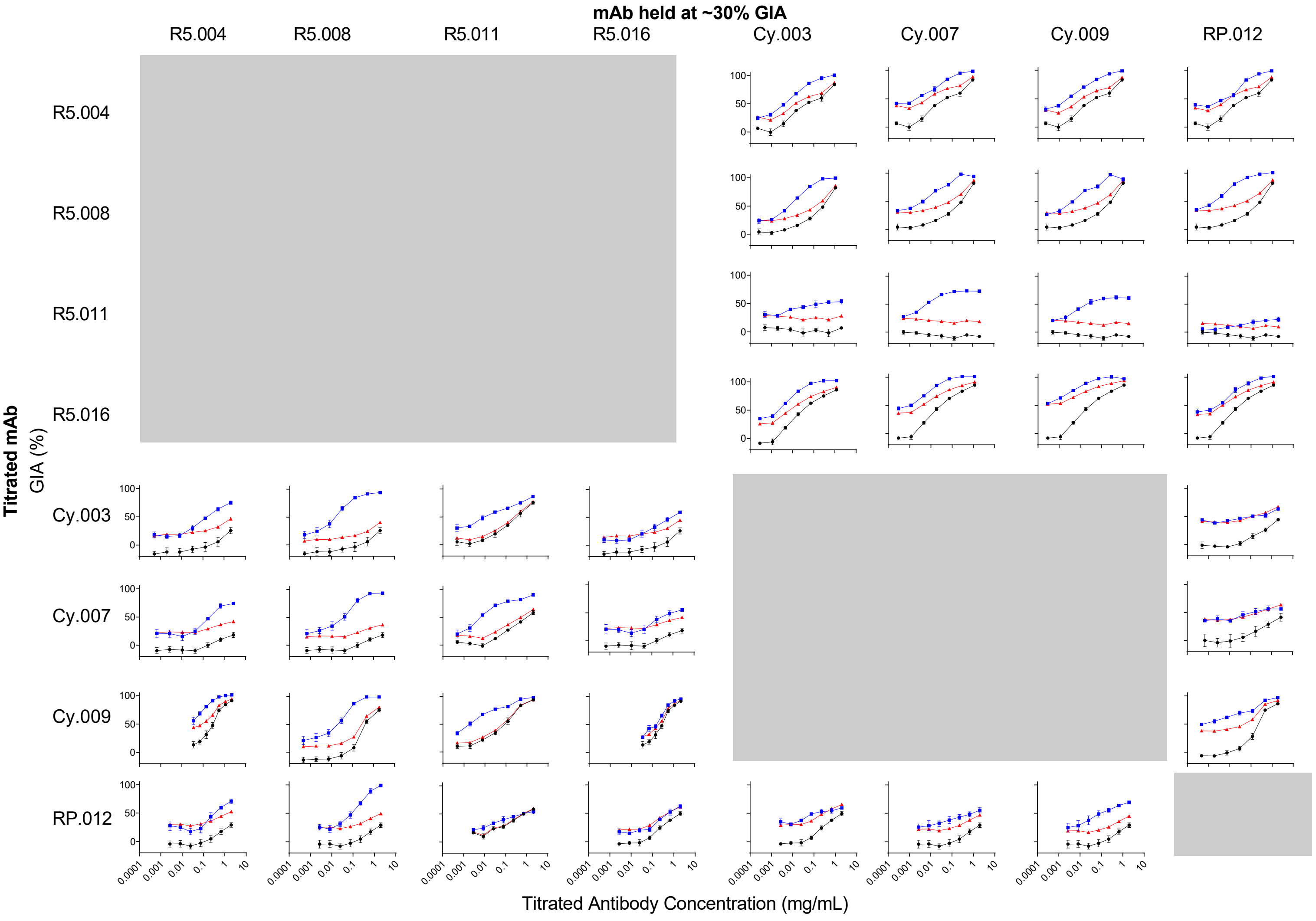
mAb held at ~30% GIA
R5.004
R5.008
R5.011
R5.016
Cy.003
Cy.007
Cy.009
RP.012
Titrated mAb
GIA (%)
100
50
0
0.0001
0.001
0.01
0.1
1
10
Titrated Antibody Concentration (mg/mL)

Figure S3

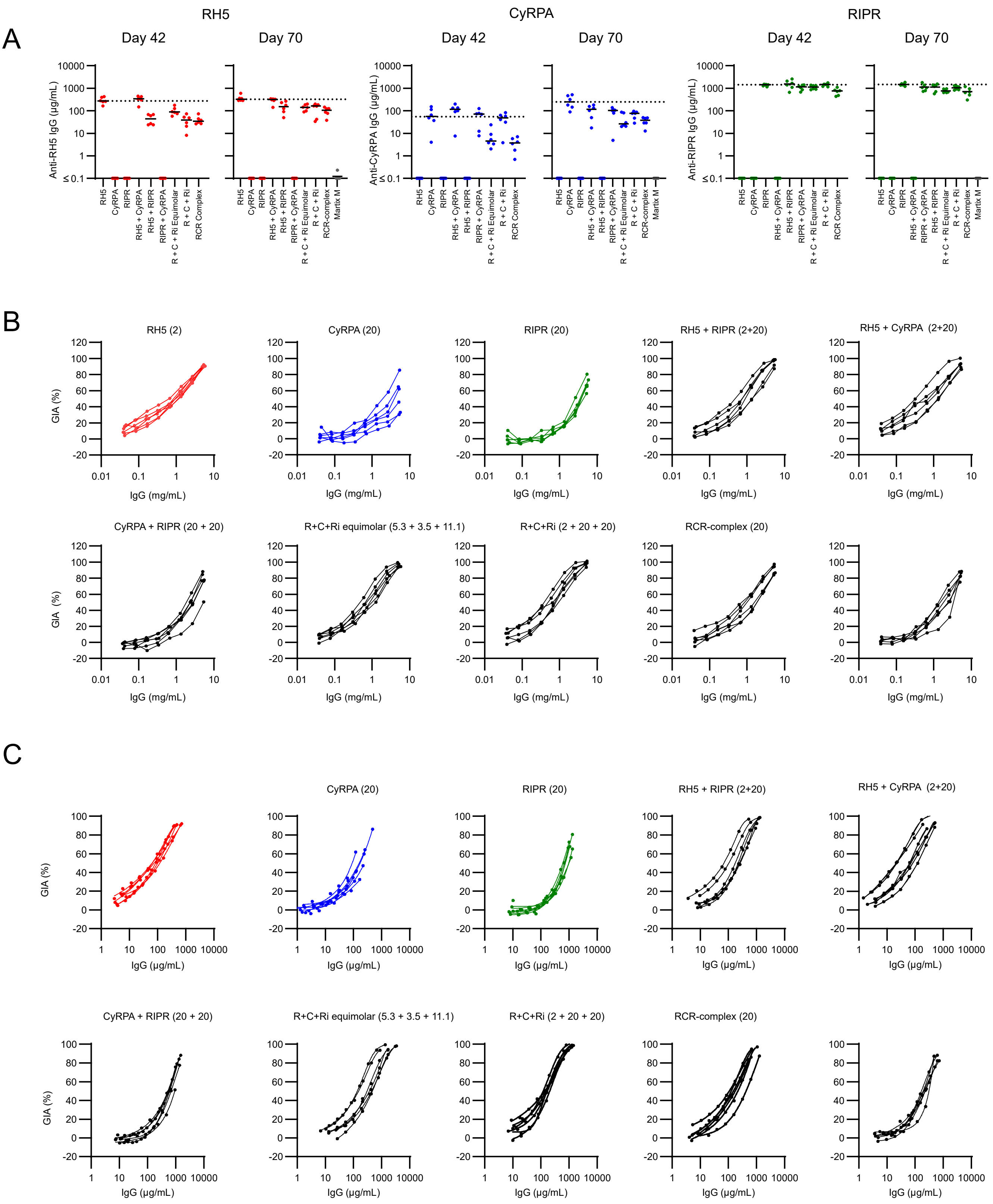
A
RH5
CyRPA
RIPR
Day 42
Day 70
Anti-RH5 IgG (µg/mL)
Anti-CyRPA IgG (µg/mL)
Anti-RIPR IgG (µg/mL)
RH5
CyRPA
RIPR
RH5 + CyRPA
RH5 + RIPR
RIPR + CyRPA
R + C + Ri Equimolar
R + C + Ri
RCR Complex
RCR-complex
Martix M
B
RH5 (2)
CyRPA (20)
RIPR (20)
RH5 + RIPR (2+20)
RH5 + CyRPA (2+20)
CyRPA + RIPR (20 + 20)
R+C+Ri equimolar (5.3 + 3.5 + 11.1)
R+C+Ri (2 + 20 + 20)
RCR-complex (20)
GIA (%)
IgG (mg/mL)
C
CyRPA (20)
RIPR (20)
RH5 + RIPR (2+20)
RH5 + CyRPA (2+20)
CyRPA + RIPR (20 + 20)
R+C+Ri equimolar (5.3 + 3.5 + 11.1)
R+C+Ri (2 + 20 + 20)
RCR-complex (20)
GIA (%)
IgG (µg/mL)

Figure S4

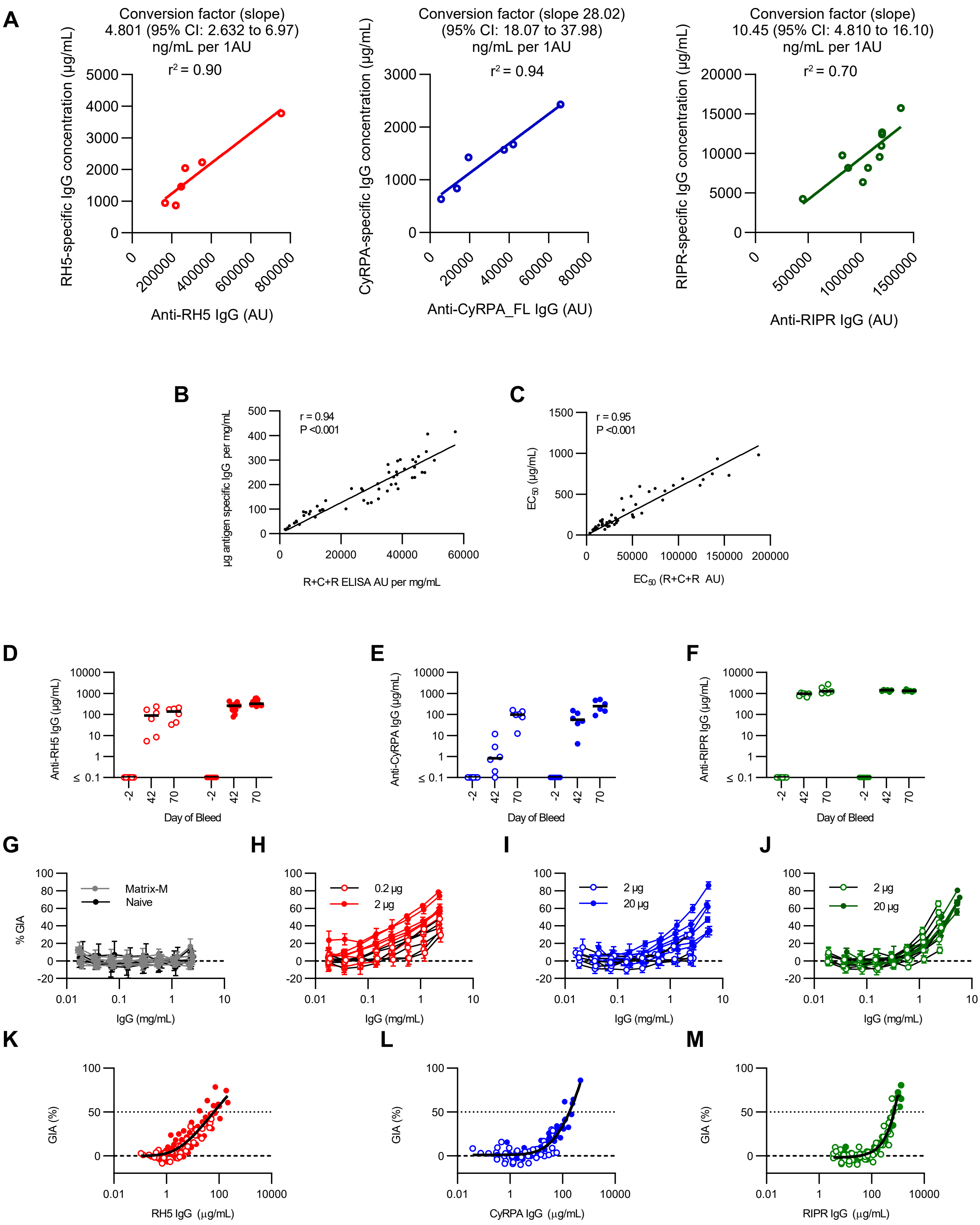


**N**

| Antigen | Dose (µg) | $EC_{50}$ (µg/mL) | 95% CI (µg/mL) |
|---|---|---|---|
| RH5 | 0.2, 2 | 64 | 50 to 89 |
| CyRPA | 2, 20 | 183 | 165 to 204 |
| RIPR | 2, 20 | 715 | 669 to 778 |

Figure S5

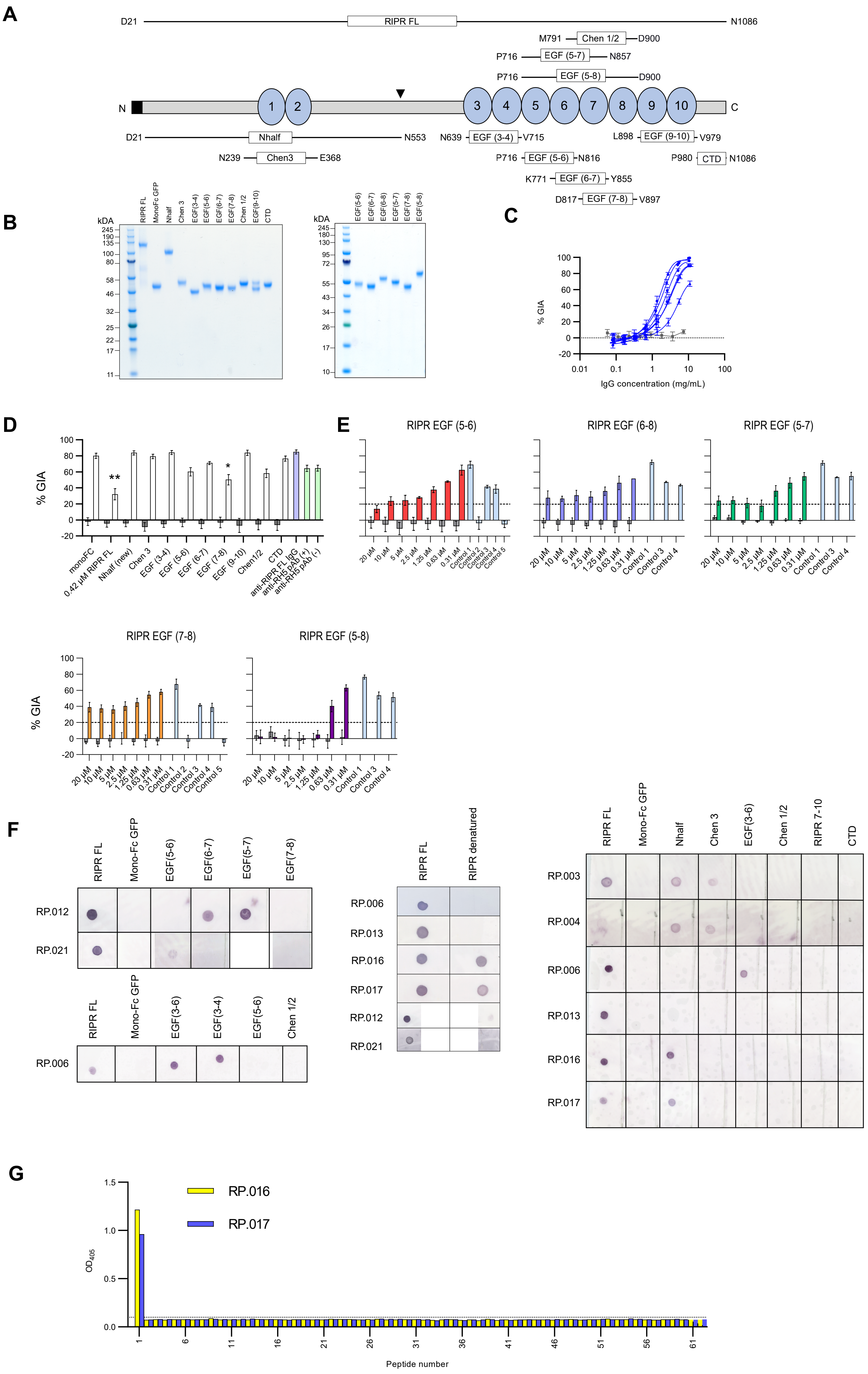

A
RIPR FL
Chen 1/2
EGF (5-7)
EGF (5-8)
Nhalf
Chen3
EGF (3-4)
EGF (9-10)
EGF (5-6)
EGF (6-7)
EGF (7-8)
CTD
B
C
% GIA
IgG concentration (mg/mL)
D
E
RIPR EGF (5-6)
RIPR EGF (6-8)
RIPR EGF (5-7)
RIPR EGF (7-8)
RIPR EGF (5-8)
F
G
RP.016
RP.017
Peptide number

Figure S6

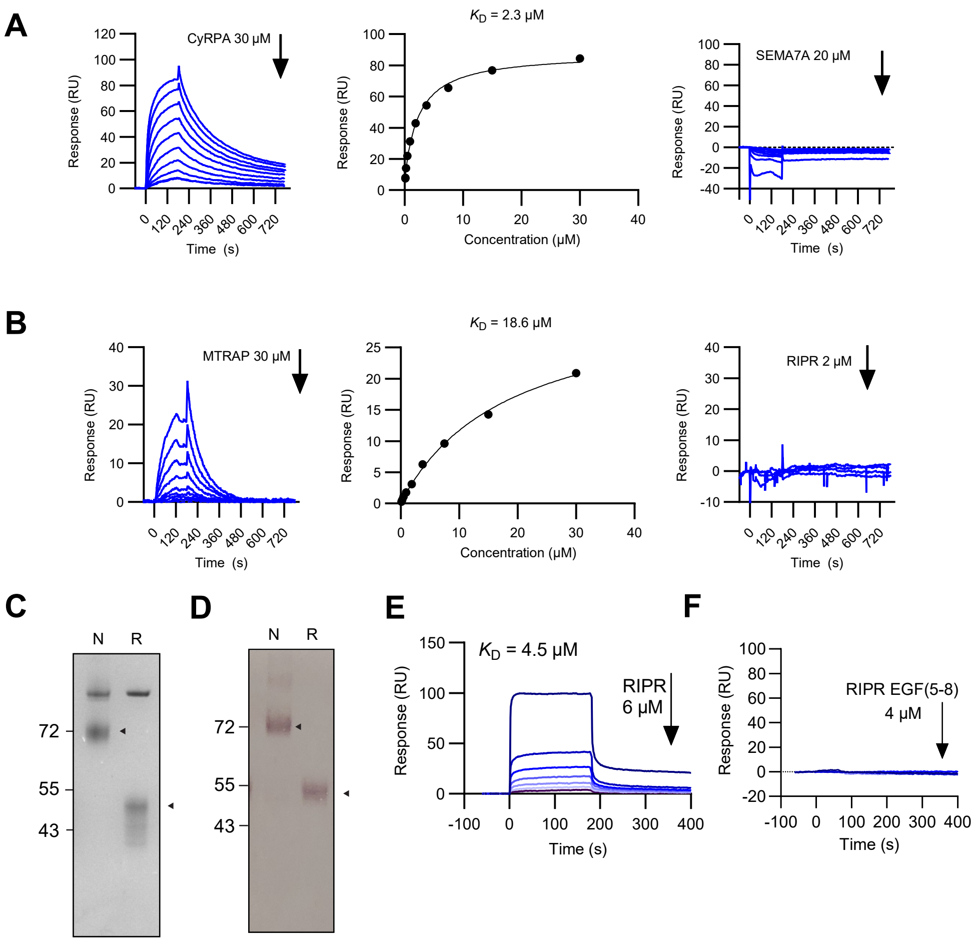
A
CyRPA 30 μM
Response (RU)
Time (s)
$K_D$ = 2.3 μM
Concentration (μM)
SEMA7A 20 μM
B
MTRAP 30 μM
$K_D$ = 18.6 μM
RIPR 2 μM
C
D
N
R
72
55
43
E
$K_D$ = 4.5 μM
RIPR
6 μM
F
RIPR EGF(5-8)
4 μM

### Figure S7

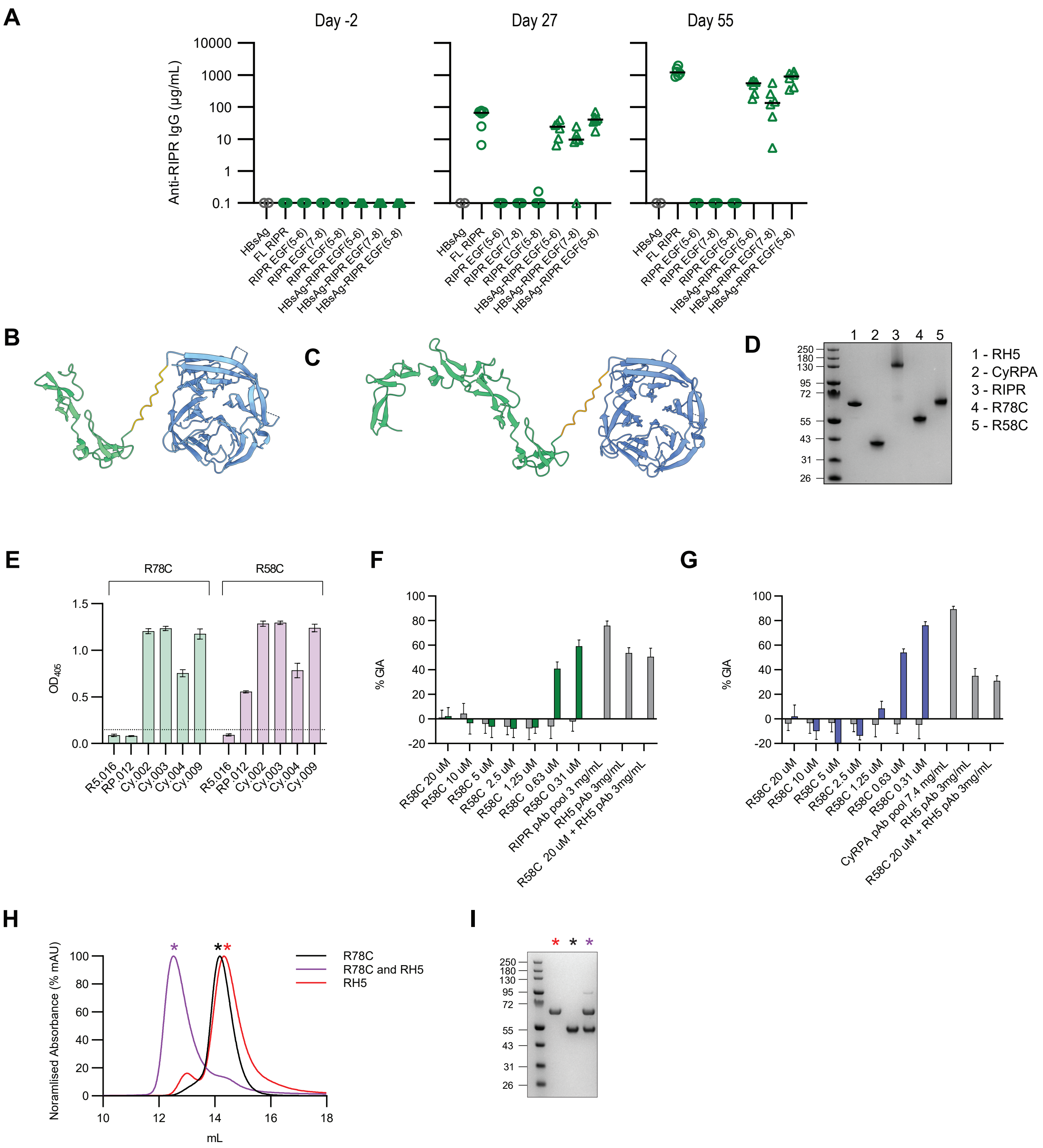

A
Day -2
Day 27
Day 55
Anti-RIPR IgG (μg/mL)
10000
1000
100
10
1
0.1
HBsAg
FL RIPR
RIPR EGF(5-6)
RIPR EGF(7-8)
RIPR EGF(5-8)
HBsAg-RIPR EGF(5-6)
HBsAg-RIPR EGF(7-8)
HBsAg-RIPR EGF(5-8)
B
C
D
1 2 3 4 5
250
180
130
95
72
55
43
31
26
1 - RH5
2 - CyRPA
3 - RIPR
4 - R78C
5 - R58C
E
R78C
R58C
OD405
1.5
1.0
0.5
0.0
R5.016
RP.012
Cy.002
Cy.003
Cy.004
Cy.009
F
% GIA
100
80
60
40
20
0
-20
R58C 20 uM
R58C 10 uM
R58C 5 uM
R58C 2.5 uM
R58C 1.25 uM
R58C 0.63 uM
R58C 0.31 uM
RIPR pAb pool 3 mg/mL
RH5 pAb 3mg/mL
R58C 20 uM + RH5 pAb 3mg/mL
G
% GIA
CyRPA pAb pool 7.4 mg/mL
RH5 pAb 3mg/mL
R58C 20 uM + RH5 pAb 3mg/mL
H
Noramlised Absorbance (% mAU)
100
80
60
40
20
0
10
12
14
16
18
mL
R78C
R78C and RH5
RH5
I
250
180
130
95
72
55
43
31
26

Figure S8

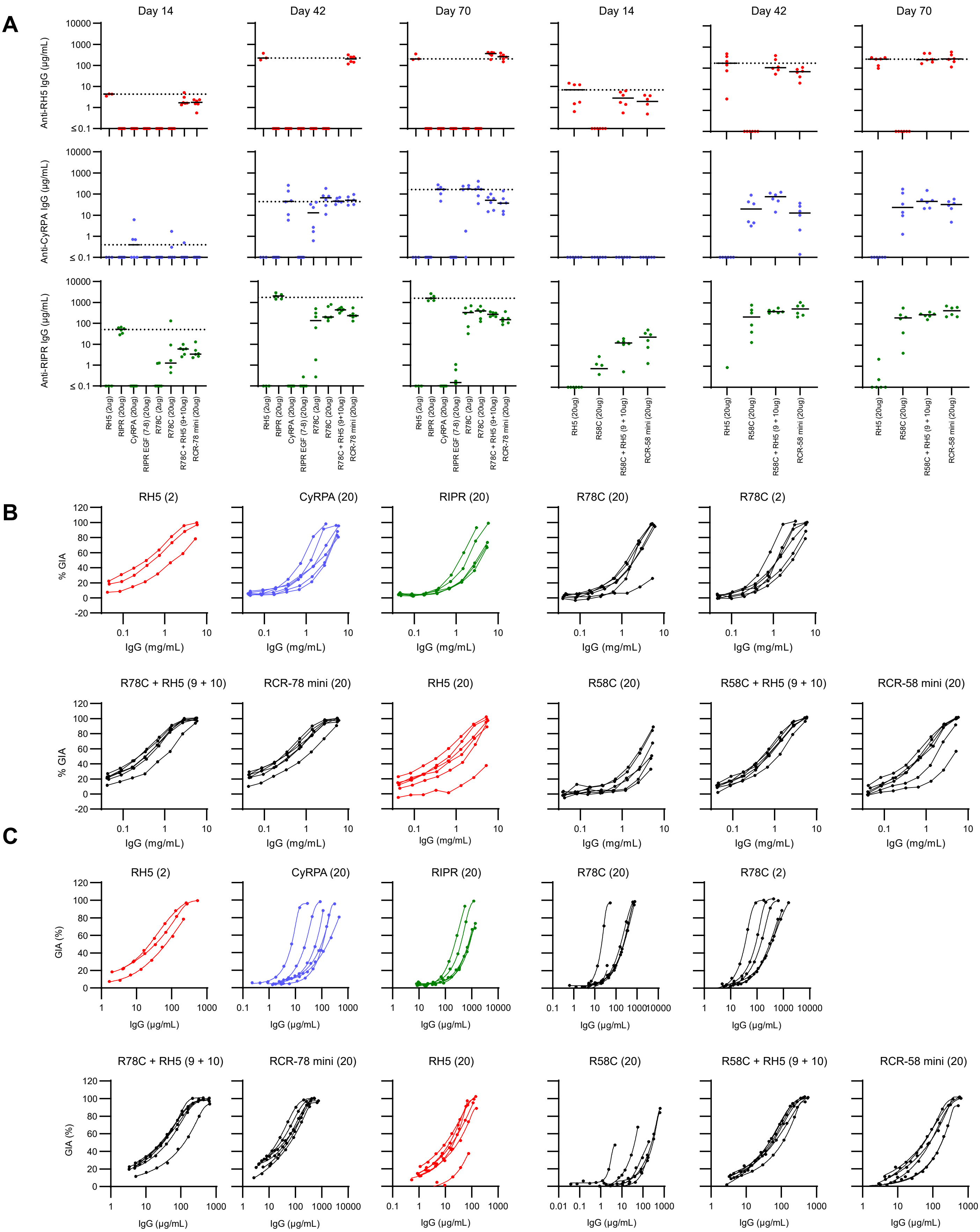

A
Day 14
Day 42
Day 70
Day 14
Day 42
Day 70
Anti-RH5 IgG (μg/mL)
Anti-CyRPA IgG (μg/mL)
Anti-RIPR IgG (μg/mL)
RH5 (2ug)
RIPR (20ug)
CyRPA (20ug)
RIPR EGF (7-8) (20ug)
R78C (2ug)
R78C (20ug)
R78C + RH5 (9+10ug)
RCR-78 mini (20ug)
RH5 (20ug)
R58C (20ug)
R58C + RH5 (9 + 10ug)
RCR-58 mini (20ug)
B
RH5 (2)
CyRPA (20)
RIPR (20)
R78C (20)
R78C (2)
R78C + RH5 (9 + 10)
RCR-78 mini (20)
RH5 (20)
R58C (20)
R58C + RH5 (9 + 10)
RCR-58 mini (20)
% GIA
IgG (mg/mL)
C
GIA (%)
IgG (μg/mL)

**Table S1 - mAb summary**

| Antigen | mAb name | % GIA at 0.5 mg/mL (3D7) | Type | Epitope bin | mAb Fc species (species origin if applicable) | Isotype used | Reference |
|---|---|---|---|---|---|---|---|
| RH5 | R5.004 | 89.99 | I | Blue | Human | IgG1 | Alanine et al. 2019 |
| | R5.013 | 65.91 | I | Blue | Human | IgG1 | Alanine et al. 2019 |
| | R5.017 | 84.31 | I | Blue | Human | IgG1 | Alanine et al. 2019 |
| | R5.019 | 86.71 | I | Blue | Human | IgG1 | Alanine et al. 2019 |
| | QA1 | 45.53 | I | Blue | Human (Mouse) | IgG1 | Douglas et al. 2014 |
| | R5.008 | 87.91 | I | Brown | Human | IgG1 | Alanine et al. 2019 |
| | R5.018 | 41.41 | I | Brown | Human | IgG1 | Alanine et al. 2019 |
| | R5.010 | 4.21 | I | Green | Human | IgG1 | Alanine et al. 2019 |
| | R5.011 | 3.37 | I | Green | Human | IgG1 | Alanine et al. 2019 |
| | R5.014 | 0.00 | I | Green | Human | IgG1 | Alanine et al. 2019 |
| | R5.002 | 12.64 | II | Orange | Human | IgG1 | Alanine et al. 2019 |
| | R5.003 | 4.50 | II | Orange | Human | IgG1 | Alanine et al. 2019 |
| | R5.006 | 13.20 | II | Orange | Human | IgG1 | Alanine et al. 2019 |
| | R5.009 | 16.00 | II | Orange | Human | IgG1 | Alanine et al. 2019 |
| | R5.015 | 5.22 | II | Orange | Human | IgG1 | Alanine et al. 2019 |
| | R5.007 | 9.68 | II | Purple | Human | IgG1 | Alanine et al. 2019 |
| | R5.016 | 85.69 | I | Red | Human | IgG1 | Alanine et al. 2019 |
| | 2AC7 | 85.54 | NT | Red | Human (Mouse) | IgG1 | Douglas et al. 2014 |
| | 9AD4 | 73.53 | I | Red | Human (Mouse) | IgG1 | Douglas et al. 2014 |
| | R5.001 | 20.19 | II | Yellow | Human | IgG1 | Alanine et al. 2019 |
| CyRPA | Cy.002 | 0.00 | II | NA | Human (Mouse) | IgG1 | Ragotte et al. 2022 |
| | Cy.003 | 28.06 | I | NA | Human (Chicken) | IgG1 | Ragotte et al. 2022 |
| | Cy.004 | 68.54 | I | NA | Human (Chicken) | IgG1 | Ragotte et al. 2022 |
| | Cy.005 | 2.68 | I | NA | Human (Mouse) | IgG1 | Ragotte et al. 2022 |
| | Cy.007 | 27.90 | I | NA | Human (Chicken) | IgG1 | Ragotte et al. 2022 |
| | Cy.009 | 74.19 | I | NA | Human (Chicken) | IgG1 | Ragotte et al. 2022 |
| | Cy.010 | 0.00 | II | NA | Human (Mouse) | IgG1 | Ragotte et al. 2022 |
| RIPR | RP.003 | 0.00 | II | EGF(1-2) | Mouse | NT | This publication |
| | RP.004 | 8.30 | II | EGF(1-2) | Mouse | NT | This publication |
| | RP.006 | 8.50 | I | EGF(3-4) | Mouse | NT | This publication |
| | RP.012 | 23.70 | I | EGF(6-7) | Mouse | IgG1 | This publication |
| | RP.013 | 1.40 | I | Unknown | Mouse | NT | This publication |
| | RP.016 | 0.00 | I | Nterm | Mouse | NT | This publication |
| | RP.017 | 1.90 | II | Nterm | Mouse | NT | This publication |
| | RP.021 | 25.80 | I | EGF(5) | Mouse | IgG2a | This publication |

**Table S2 - Summary statistics for Figure 4.** One-way ANOVA on log-transformed data with Dunnett's multiple comparison test

| Panel A | | |
|---|---|---|
| **Test** | **Summary** | **Adjusted P value** |
| RH5 vs. RH5 + CyRPA | ns | 0.2311 |
| RH5 vs. RH5 + RIPR | **** | <0.0001 |
| RH5 vs. R + C + Ri Equimolar | **** | <0.0001 |
| RH5 vs. R + C + Ri | **** | <0.0001 |
| RH5 vs. RCR-complex | **** | <0.0001 |
| **Panel B** | | |
| CyRPA vs. RH5 + CyRPA | *** | 0.0002 |
| CyRPA vs. RIPR + CyRPA | **** | <0.0001 |
| CyRPA vs. R + C + Ri Equimolar | **** | <0.0001 |
| CyRPA vs. R + C + Ri | **** | <0.0001 |
| CyRPA vs. RCR-complex | **** | <0.0001 |
| **Panel C** | | |
| RIPR vs. RIPR + CyRPA | ns | 0.0833 |
| RIPR vs. RH5 + RIPR | ns | 0.0819 |
| RIPR vs. R + C + Ri Equimolar | **** | <0.0001 |
| RIPR vs. R + C + Ri | * | 0.0187 |
| RIPR vs. RCR-complex | **** | <0.0001 |

A

B

C

**Table S3 - Summary statistics for Figure 6.** One-way ANOVA on log-transformed data with Dunnett's multiple comparison test

**A**

Panel A

| Test | Summary | Adjusted P Value |
|---|---|---|
| RH5 (2ug) vs. CyRPA (20ug) | **** | <0.0001 |
| RH5 (2ug) vs. RIPR (20ug) | **** | <0.0001 |
| RH5 (2ug) vs. RIPR EGF (7-8)(20ug) | **** | <0.0001 |
| RH5 (2ug) vs. R78C (2ug) | **** | <0.0001 |
| RH5 (2ug) vs. R78C (20ug) | **** | <0.0001 |
| RH5 (2ug) vs. R78C + RH5 (9+10ug) | ns | 0.0534 |
| RH5 (2ug) vs. RCR-78 mini (20ug) | ns | 0.9971 |

Panel B

| Test | Summary | Adjusted P Value |
|---|---|---|
| RH5 (20ug) vs. R58C (20ug) | **** | <0.0001 |
| RH5 (20ug) vs. R58C + RH5 (9+10ug) | ns | 0.5648 |
| RH5 (20ug) vs. RCR-58 mini (20ug) | ns | 0.6526 |

**B**

Panel C

| Test | Summary | Adjusted P Value |
|---|---|---|
| CyRPA (20ug) vs. RH5 (2ug) | **** | <0.0001 |
| CyRPA (20ug) vs. RIPR (20ug) | **** | <0.0001 |
| CyRPA (20ug) vs. RIPR EGF (7-8)(20ug) | **** | <0.0001 |
| CyRPA (20ug) vs. R78C (2ug) | ns | 0.1818 |
| CyRPA (20ug) vs. R78C (20ug) | ns | >0.9999 |
| CyRPA (20ug) vs. R78C + RH5 (9+10ug) | ns | 0.4637 |
| CyRPA (20ug) vs. RCR-78 mini (20ug) | ns | 0.3631 |

Panel D

| Test | Summary | Adjusted P Value |
|---|---|---|
| CyRPA vs. RH5 (20ug)† | **** | <0.0001 |
| CyRPA vs. R58C (20ug)† | * | 0.0113 |
| CyRPA vs. R58C + RH5 (9+10ug)‡ | ns | 0.1636 |
| CyRPA vs. RCR-58 mini (20ug)† | * | 0.0208 |

**C**

Panel E

| Test | Summary | Adjusted P Value |
|---|---|---|
| RIPR (20ug) vs. RH5 (2ug) | **** | <0.0001 |
| RIPR (20ug) vs. CyRPA (20ug) | **** | <0.0001 |
| RIPR (20ug) vs. RIPR EGF (7-8) (20ug) | **** | <0.0001 |
| RIPR (20ug) vs. R78C (2ug) | ** | 0.0012 |
| RIPR (20ug) vs. R78C (20ug) | * | 0.0103 |
| RIPR (20ug) vs. R78C + RH5 (9+10ug) | ** | 0.0045 |
| RIPR (20ug) vs. RCR-78 mini (20ug) | *** | 0.0002 |

Panel F

| Test | Summary | Adjusted P Value |
|---|---|---|
| RIPR vs. RH5 (20ug)‡ | **** | <0.0001 |
| RIPR vs. R58C (20ug)‡ | *** | 0.0005 |
| RIPR vs. R58C + RH5 (9+10ug)‡ | * | 0.0224 |
| RIPR vs. RCR-58 mini (20ug)‡ | ns | 0.088 |

†Performed on CyRPA data from panel C

‡ Performed on RIPR data from panel E

**Table S4 – Details of RIPR truncations produced.**

| RIPR fragment description | Amino acids | Expression in Expi293 cells | MS/MS verified | Intact mass verified |
|---|---|---|---|---|
| monoFc + GFP | N/A | ✓ | ✓ | ✓ |
| monoFc + N-half | D21-N553 | ✓ | ✓ | ✗ |
| monoFc + Chen3 | N239-E368 | ✓ | ✓ | ✓ |
| monoFc + EGF domains 3-6 | N639-N816 | ✓ | ✓ | ✓ |
| monoFc + Chen1/2 | N791-D900 | ✓ | ✓ | ✓ |
| monoFc + EGF domains 7-10 | D817-V979 | ✓ | ✓ | ✓ |
| monoFc + EGF domains 3-4 | N639-V715 | ✓ | ✓ | ✓ |
| monoFc + EGF domains 5-6 | V716-N816 | ✓ | ✓ | ✓ |
| monoFc + EGF domains 6-7 | K771-Y855 | ✓ | ✓ | ✓ |
| monoFc + EGF domains 7-8 | D817-V897 | ✓ | ✓ | ✓ |
| monoFc + EGF domains 9-10 | L898-V979 | ✓ | ✓ | ✓ |
| monoFc + EGF domains 5-8 | V716-D900 | ✓ | ✓ | ✓ |
| monoFc + EGF domains 5-7 | V716-N857 | ✓ | ✓ | ✓ |
| monoFc + EGF domains 3-6 | K771 - N816 | Poor | ✓ | ✗ |
| monoFc + EGF domains 7-10 | D817 - V979 | Poor | ✓ | ✗ |
| monoFc + C-terminal domain | P980-N1086 | ✓ | ✓ | ✓ |

**Table S5 – Details of PCR primers used to make RIPR truncations.**

| Name | Sequence | Target | Amino Acid coodinate | Enzyme sites | Orientation |
|---|---|---|---|---|---|
| NhalfCoFW | gatcggatccGATCTGATCGAGGGCATCTT | Codon optimised (3133) Nhalf (NH) | 21 | 4 Bases-BamHI-Ripr | FW |
| NhalfCoRV | gatcggtaccaccGTTAGAGTTGGTGTGGGGGT | Codon optimised (3133) Nhalf (NH) | 533 | 4 Bases-KpnI-G-Ripr | RV |
| BGWP007 | gatcggatccAATATCAACAACATGAATAT | CO PfRipr (5285) Chalf region (CH) | 534 | 4 Bases-BamHI-Ripr | FW |
| BGWP008 | gatcggtaccaccGTTCTGGTTGGAGTAATAGA | CO PfRipr (5285) Chalf region (CH) | 1086 | 4 Bases-KpnI-G-Ripr | RV |
| BGWP009 | gatcggatccCGGTGCACCCAGGACATCTG | CO PfRipr (5285) EGF1-2 (E12D) | 287 | 4 Bases-BamHI-Ripr | FW |
| BGWP010 | gatcggtaccaccGTAGCACTTGTTGTTGTGGT | CO PfRipr (5285) EGF1-2 (E12D) | 362 | 4 Bases-KpnI-G-Ripr | RV |
| BGWP013 | gatcggatccAACCTGTGCAACAAGTGTCA | CO PfRipr (5285) EGF3-10 (E310D) | 639 | 4 Bases-BamHI-Ripr | FW |
| BGWP014 | gatcggtaccaccCACGCAGCTGTCGTCCTTGT | CO PfRipr (5285) EGF3-10 (E310D) | 979 | 4 Bases-KpnI-G-Ripr | RV |
| BGWP015 | gatcggatccCCCAACACCAACGAGTACGACGA | CO PfRipr (5285) CTD | 980 | 4 Bases-BamHI-Ripr | FW |
| BGWP019 | gatcggatccAACATCCAGTACCAGTGCAT | CO PfRipr (5285) Chen3 | 238 | 4 Bases-BamHI-Ripr | FW |
| BGWP020 | gatcggatccAACGAAGAGACAGACATCGT | CO PfRipr (5285) Chen1/2 | 791 | 4 Bases-BamHI-Ripr | FW |
| BGWP021 | gatcggtaccaccATCTTCCAGCACGCACTTCC | CO PfRipr (5285) Chen1/2 | 900 | 4 Bases-KpnI-G-Ripr | RV |
| BGWP022 | gatcggtaccaccATTCAGGATGCACTCGCCTC | CO PfRipr (5285) EGF3-6 (E36D) | 816 | 4 Bases-KpnI-G-Ripr | RV |
| BGWP023 | gatcggatccGACTACTGCAAGGACATCAA | CO PfRipr (5285) EGF 7-10 (E710D) | 817 | 4 Bases-BamHI-Ripr | FW |
| BGWP042 | gatcggtaccaccCACGCATTTGCCCTTCACGT | CO PfRipr (5285) E34D | 716 | 4 Bases-KpnI-G-Ripr | RV |
| BGWP043 | gatcggatccCCCGATAACAAGTGCGACCT | CO PfRipr (5285) E56D | 716 | 4 Bases-BamHI-Ripr | FW |
| BGWP044 | gatcggtaccaccCACGCACTTCCCGTTCACGG | CO PfRipr (5285) E78D | 897 | 4 Bases-KpnI-G-Ripr | RV |
| BGWP045 | gatcggatccCTGGAAGATAAGTGCGTGCA | CO PfRipr (5285) E910D | 898 | 4 Bases-BamHI-Ripr | FW |
| BGWP054 | gatcggtaccaccGTTTTTGGCAATGCAGTTGA | CO PfRipr (5285) EGF 5 | 770 | 4 Bases-KpnI-G-Ripr | RV |
| BGWP055 | gatcggatccAAGTGCAAGCGGAAAGAGTA | CO PfRipr (5285)EGF 6 | 771 | 4 Bases-BamHI-Ripr | FW |
| BGWP056 | gatcggtaccaccATAGATGCACTCCCCCTTGT | CO PfRipr (5285) EGF 7 | 855 | 4 Bases-KpnI-G-Ripr | RV |
| BGWP057 | gatcggatccGAGAACAGCTGCCTGATCAA | CO PfRipr (5285) EGF 8 | 856 | 4 Bases-BamHI-Ripr | FW |
| BGWP068 | gatcggatccGCCAAAAACAAGTGCAAGCG | CO PfRipr (5285) EGF 6 | 768 | 4 Bases-BamHI-Ripr | FW |
| BGWP069 | gatcggtaccaccGTCATTCAGGATGCACTCGC | CO PfRipr (5285) EGF 6 | 817 | 4 Bases-KpnI-G-Ripr | RV |
| BGWP070 | gatcggatccCTGAATGACTACTGCAAGGA | CO PfRipr (5285) EGF 7 | 815 | 4 Bases-BamHI-Ripr | FW |
| BGWP071v2 | gatcggtaccaccGTTCTCATAGATGCACTCCC | CO PfRipr (5285) EGF 7 | 857 | 4 Bases-KpnI-G-Ripr | RV |
| BGWP072 | gatcggatccTATGAGAACAGCTGCCTGAT | CO PfRipr (5285) EGF 8 | 855 | 4 Bases-BamHI-Ripr | FW |
| BGWP073 | gatcggtaccaccATCTTCCAGCACGCACTTCC | CO PfRipr (5285) EGF 8 | 900 | 4 Bases-KpnI-G-Ripr | RV |
| BGWP074 | gatcggatccCTGGAAGATAAGTGCGTGCA | CO PfRipr (5285) EGF 9 | 989 | 4 Bases-BamHI-Ripr | FW |
| BGWP075 | gatcggtaccaccATTCTGGATCAGGCACACGC | CO PfRipr (5285) EGF 9 | 941 | 4 Bases-KpnI-G-Ripr | RV |
| BGWP076 | gatcggtaccaccGGTGAAGCTCTCGTCGTACT | CO PfRipr (5285) EGF 10 | 990 | 4 Bases-KpnI-G-Ripr | RV |

**Table S6 – Amino acid sequences of R78C and R58C**

| Protein | Amino acid sequence (signal peptide underlined). |
|---|---|
| R78C | MGWSCIILFLVATATGVHSDYCKDINCKENEECSIVNFKPECVCKENLKKNNKGECIYENSCLINEGNCPKDSKCIYRE<br>YKPHECVCNKQGHVAVNGKCVGGGGSGGGGSAHIVMVDAYKPTKGGGGSGGGGSDSRHVFIRTELSFIKNNVPCI<br>RDMFFIYKRELYNICLDDLKGEEDETHIYVQKKVKDSWITLNDLFKETDLTGRPHIFAYVDVEEIIILLCEDEEFSNRKKD<br>MTCHRFYSNDGKEYNNAEITISDYILKDKLLSSYVSLPLKIENREYFLICGVSPYKFKDDNKKDDILCMASHDKGETWG<br>TKIVIKYDNYKLGVQYFFLRPYISKNDLSFHFYVGDNINNVKNVNFIECTHEKDLEFVCSNRDFLKDNKVLQDVSTLND<br>EYIVSYGNDNNFAECYIFFNNENSILIKPEKYGNTAAGCYGGTFVKIDENRALFIYSSSQGIYNIHTIYYANYEGGGGS E<br>PEA |
| R58C | MGWSCIILFLVATATGVHSPDNKCDLSCPSNKVCVIENGKQTCKCSERFVLENGVCICANDYKMEDGINCIAKNKCK<br>RKEYENICTNPNEMCAYNEETDIVKCECKEHYYRSSRGECILNDYCKDINCKENEECSIVNFKPECVCKENLKKNNKGE<br>CIYENSCLINEGNCPKDSKCIYREYKPHECVCNKQGHVAVNGKCVLEDGGGGSGGGGSAHIVMVDAYKPTKGGGG<br>SGGGGSDSRHVFIRTELSFIKNNVPCIRDMFFIYKRELYNICLDDLKGEEDETHIYVQKKVKDSWITLNDLFKETDLTGR<br>PHIFAYVDVEEIIILLCEDEEFSNRKKDMTCHRFYSNDGKEYNNAEITISDYILKDKLLSSYVSLPLKIENREYFLICGVSPY<br>KFKDDNKKDDILCMASHDKGETWGTKIVIKYDNYKLGVQYFFLRPYISKNDLSFHFYVGDNINNVKNVNFIECTHEKD<br>LEFVCSNRDFLKDNKVLQDVSTLNDEYIVSYGNDNNFAECYIFFNNENSILIKPEKYGNTAAGCYGGTFVKIDENRALFI<br>YSSSQGIYNIHTIYYANYEGGGGSEPEA |
